## Supporting Information for "Intramembrane cleavage of TREM2 is determined by its intrinsic structural dynamics"

**Supporting Information Figures**

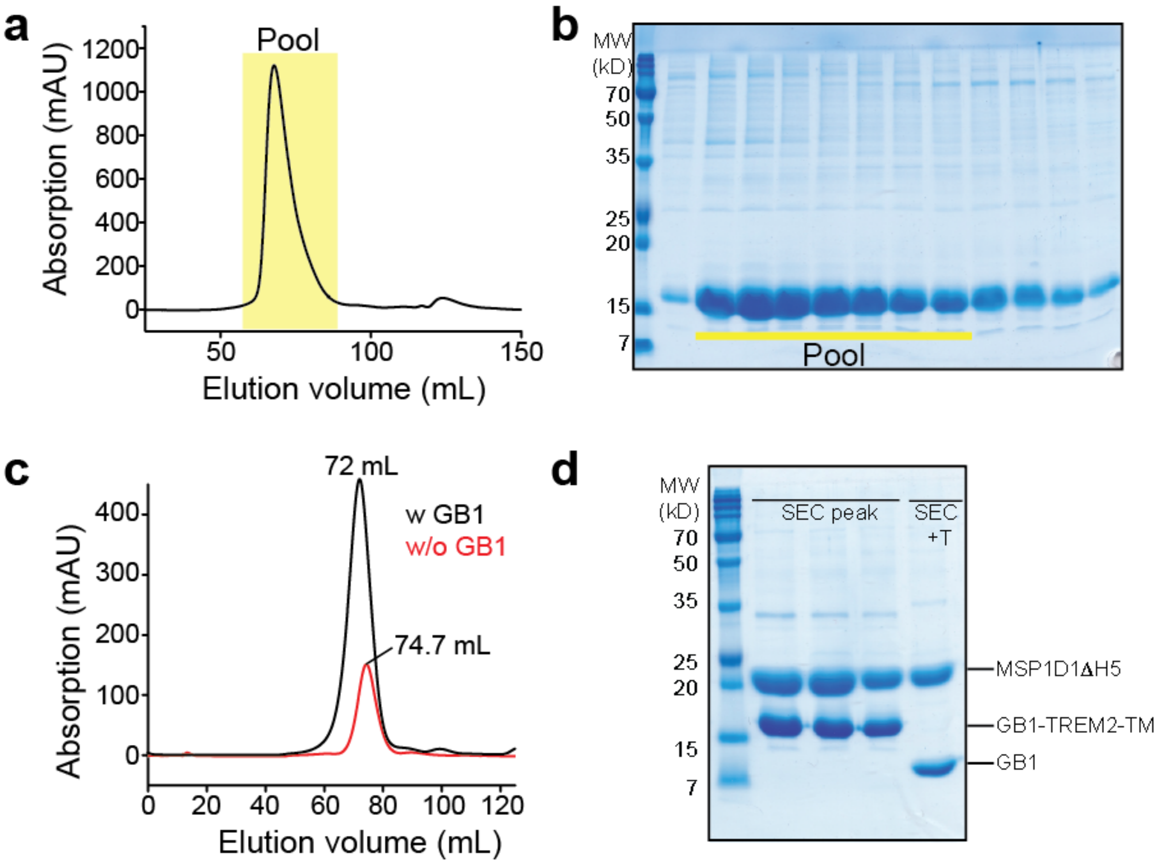


**Fig. S1. Purification of TREM2-TMH in DPC micelles.** (a) Size exclusion chromatogram (SEC) of TREM2-TMH in DPC micelles using a Superdex200 column (124 mL bed volume). The indicated region of the chromatogram was pooled and used for NMR investigations. (b) SDS-PAGE of SEC fractions containing TREM2-TMH. (c) SEC chromatogram of MSP1D1ΔH5 nanodisc-incorporated TREM2-TMH fused to GB1 (black line) or after its removal by thrombin protease cleavage, followed by Ni-NTA chromatography (red line). The apparent increase in elution volume for the cleaved protein without GB1 indicates a reduction in size. (d) SDS-PAGE of TREM2-TMH in nanodiscs before and after addition of thrombin protease for GB1 removal. T: thrombin. Due to its small size of ~5kDa, TREM2-TMH is not visible on the gel. Co-elution of TREM2-TMH and the membrane scaffold protein (MSP) in the same SEC fractions indicates successful nanodisc incorporation.

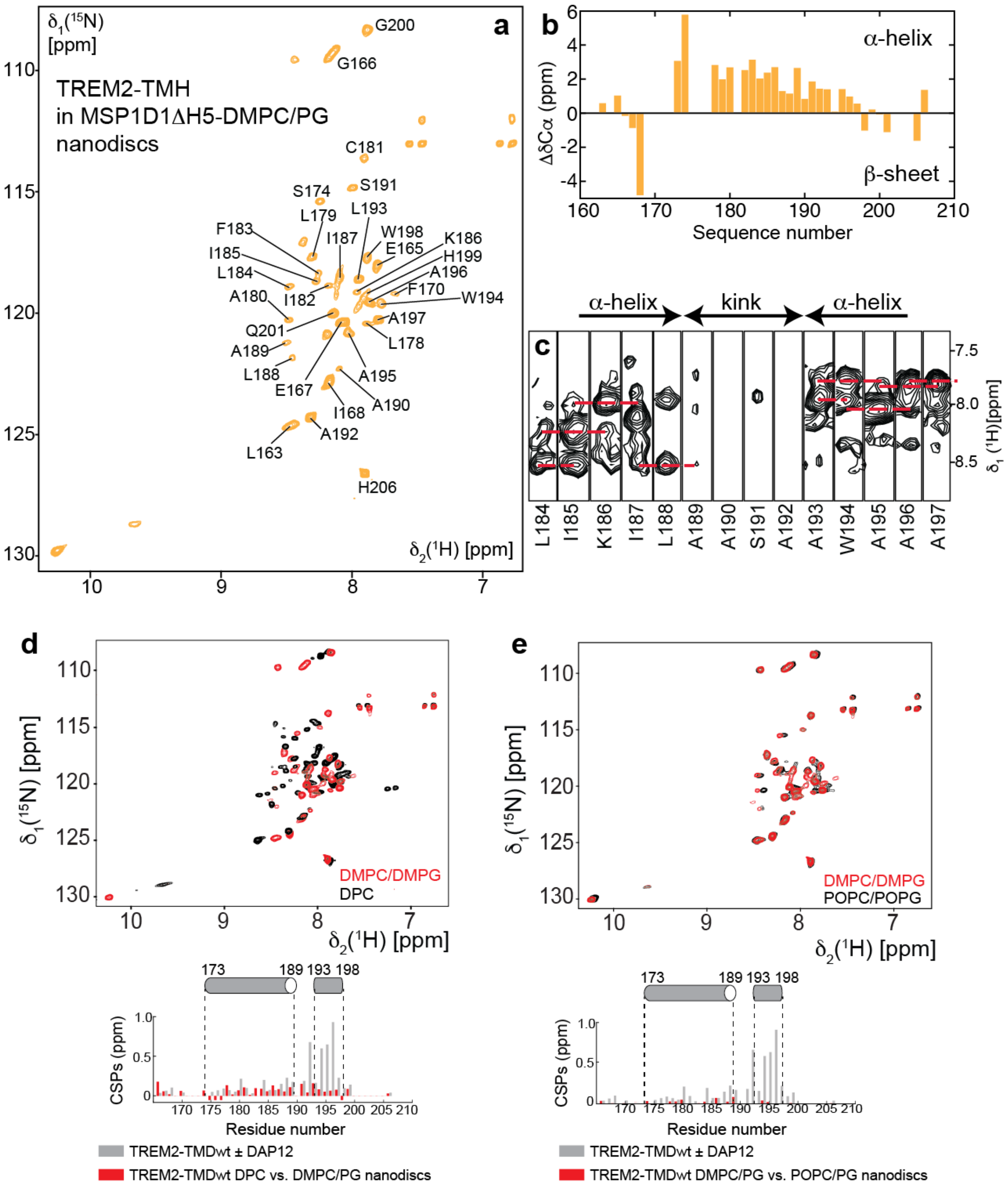


**Fig. S2. NMR structural investigations on TREM2-TMH in phospholipid nanodiscs.** (a) 2D-[$^{1}$H,$^{15}$N]-TROSY spectrum at 37°C of *U*-$^{2}$H,$^{13}$C,$^{15}$N-labeled TREM2-TMH in nanodiscs composed of MSP1D1ΔH5 and DMPC/DMPG (3:1) lipids. Assigned backbone amide resonances are labeled, as obtained with 3D-HNCA and 3D-NOESY spectra. (b) Cα secondary chemical shifts of TREM2-TMH in nanodiscs indicate α-helical secondary structure (positive values) between residues 173 and 198. Cβ chemical shift information could not be obtained for this sample due to sensitivity issues. No data could be obtained for proline stretches and regions where no signal could be observed in the TROSY spectrum. (c) 3D-$^{15}$Nedited-[$^{1}$H,$^{1}$H]-NOESY spectra show that the transmembrane helical conformation is interrupted by an unstructured stretch between residues 189 to 192, as indicated by a lack of sequential contacts between amide protons (red broken lines). (d) Overlay of 2D-[$^{1}$H,$^{15}$N]-TROSY spectra of TREM2-TMH in DPC micelles (black) and DMPC/DMPG (3:1) nanodiscs. The chemical shift perturbations (CSPs) of TREM2-TMH in DPC *versus* nanodiscs are plotted below the spectrum in red. (e) Overlay of 2D-[$^{1}$H,$^{15}$N]-TROSY spectra of TREM2-TMH in POPC/POPG (3:1) (black) and DMPC/DMPG (3:1) (red) nanodiscs. The CSPs of TREM2-TMH in DMPC/PG *versus* POPC/PG nanodiscs are plotted below the spectrum in red. Grey bars in (d) and (e) are CSP values in DPC micelles upon the addition of DAP12 for comparison.

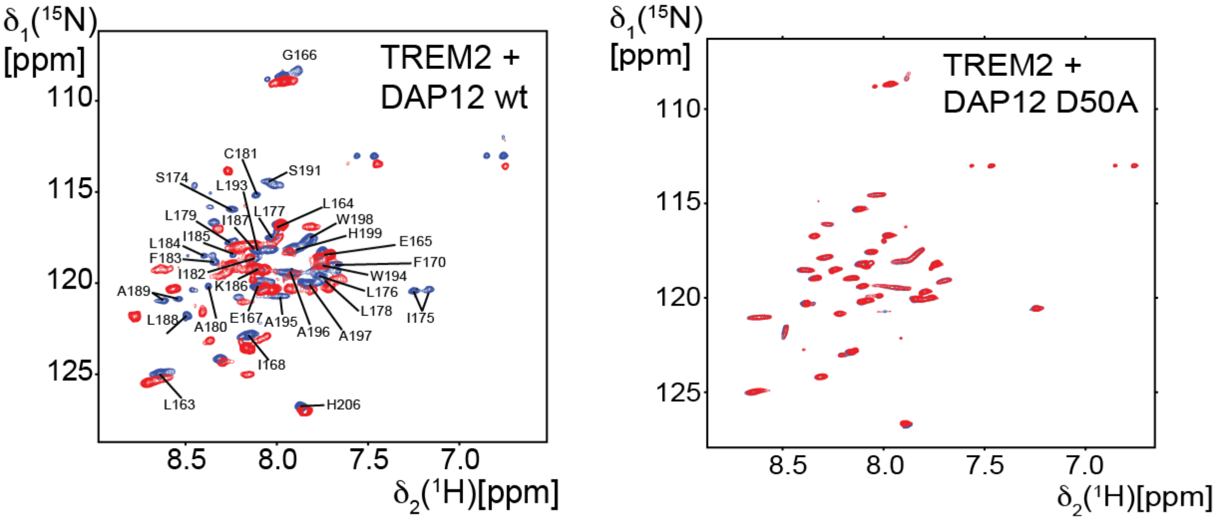


**Fig. S3. Interaction between TREM2-TM and DAP12 probed by 2D-TROSY spectra.** TREM2 interacts with DAP12 wild-type as can be seen by pronounced chemical shift perturbations (left). In contrast, no chemical shift changes are observed if the DAP12 D50A variant is added to *U*-$^{2}$H,$^{15}$N-labeled TREM2-TM (right). Blue spectrum: TREM2-TM, red spectrum: TREM2-TMH in complex with either DAP12-wt or the D50A variant.

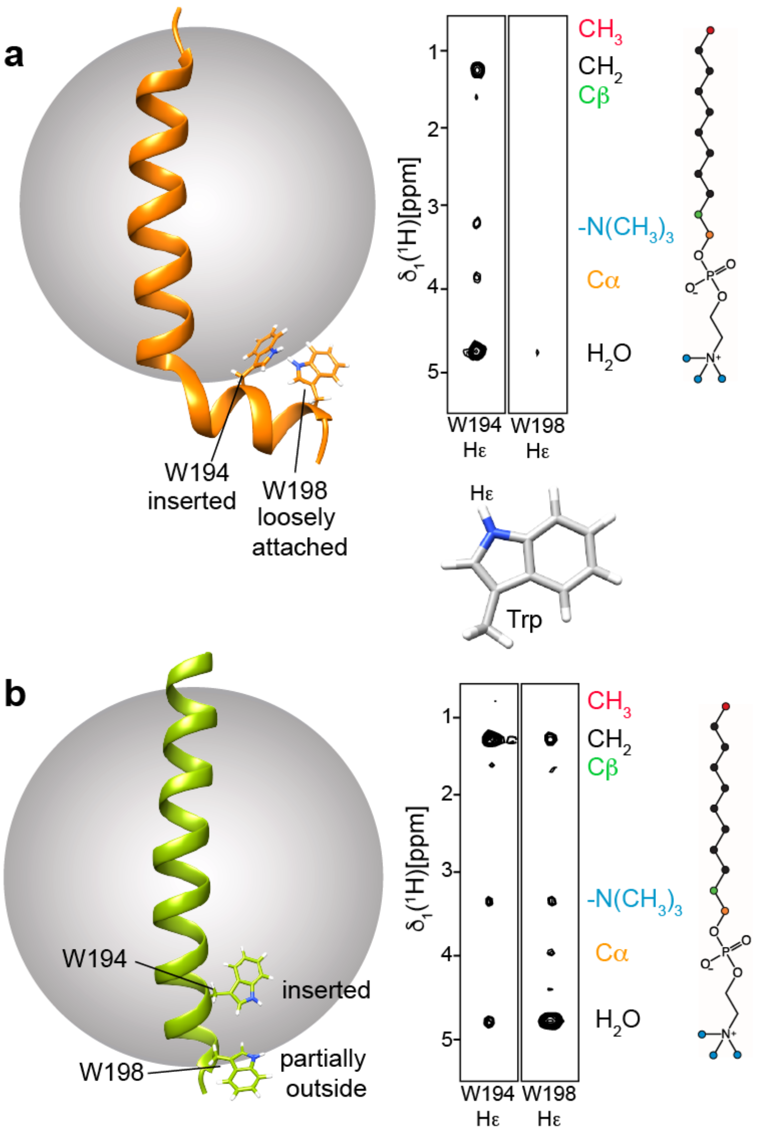


**Fig. S4. Insertion of tryptophan side chains into the hydrophobic membrane region.** (a) The Hε protons of the two tryptophan (W) residues in TREM2-TMH wt show distinct NOE patterns with the individual regions of the DPC molecule. For W194, strong NOE cross-peaks to the DPC methylene moieties can be observed, indicative of a membrane location of this side chain. Contacts to the cholate group of DPC and the solvent are visible, too, suggesting a peripheral location. For W198, no signal to any part of DPC can be observed, indicative of a dynamic state outside the micelle. (b) For the TREM2-TMH K186A variant, strong contacts for both Trp residues to the inside of the micelle can be observed. Since W198 is located at the end of the TMH, its higher solvent accessibility gives rise to a stronger NOE signal to $H_2O$ at 4.7 ppm.
